# Abundance, fatty acid composition and saturation index of neutral lipids are significantly different between isogenic primary and metastatic colon cancer cell lines

**DOI:** 10.1101/690685

**Authors:** Rimsha Munir, Jan Lisec, Carsten Jaeger, Nousheen Zaidi

## Abstract

Lipid droplets, the dynamic organelles that store triglycerides (TG) and cholesterol esters (CE), are highly accumulated in colon cancer cells. This work studies the TG and CE subspecies profile in colon carcinoma cells lines, SW480 derived from primary tumor, and SW620 derived from a metastasis of the same tumor. It was previously reported that the total TG and CE content is dramatically higher in SW620 cells; however, TG and CE subspecies profile has not been investigated in detail. The presented work confirms that total TG and CE content is significantly higher in SW620 cells. Moreover, the FA-composition of TGs is significantly altered in SW620 cells, with significant decrease in the abundance of saturated triglycerides. This resulted in significantly decreased TG saturation index in SW620 cells. The saturation index of CEs was also significantly decreased in SW620 cells. The saturation indices of some other major lipid classes were either similar or only moderately different between SW480 and SW620 cells. We also compared the expression of metabolic genes that may regulate these changes in the lipidomic profiles.

## Introduction

Lipidomic landscape of colorectal cancer (CRC) cells is significantly modified in comparison to that in their non-malignant counterparts (reviewed in [1]). Primary and metastatic colon cancer cells also display differences in their lipidomic profiles [2]. Previous works have taken advantage of SW480/SW620 cell line pair, which represents an accepted model to study metastatic progression [3]. SW480 is derived from a primary tumor, and SW620 is derived from a metastasis of the same tumor [4]. Multiple differences in the lipidomic profiles of these two cell lines have been noted [2]. For instance; increased levels of plasmanylcholine, while decreased levels of plasmenylethanolamine, C-16 containing sphingomyelin and ceramide were observed [2]. It has been shown that, SW620 cells display dramatic increase in total triglyceride (TG) and cholesterol ester (CE) levels [2]. However, the TG and CE subspecies profiles has not been explored in detail. Studying the TG and CE profile CRC cells is particularly important because these neutral lipids from hydrophobic core of lipid droplets (LDs) –that are highly accumulated in CRC cells [5]. Previous works have shown that exposure to excessive poly unsaturated fatty acids (PUFAs) induces LD formation in both SW480 and SW620 cells [6]. Interestingly, SW620 cells accumulate PUFA-enriched CEs, while SW480 cells accumulates PUFA-enriched TGs [6]. This indicates differential regulation of TG and CE content in SW480 and SW620 cells. Herein, we study the differences in TG- and CE-subspecies profiles and saturation index (SI) between SW480 and SW620 cells. Moreover, the expression of various metabolic genes that may bring about these lipidomic changes were also analyzed.

## Materials and Methods

### Cell culture

The SW480 and SW620, cell lines were purchased from American Type Culture Collection and were maintained in DMEM (Gibco, 31966-021) supplemented with 10% fetal bovine serum (FBS) (Sigma, F75240) and penicillin-streptomycin solution (Corning, 30-002-CI). Cell cultures were maintained in the atmosphere of 5% CO2 and 37°C. Cell lines were commercially authenticated (Eurofins, Germany) and mycoplasma tested prior to submission of this manuscript.

### Gene Expression Analysis

For quantitative RT-PCR, total RNA was extracted from cell pellets using Quick-RNA™ MiniPrep Plus (Zymo Research). All RNA samples were reverse-transcribed into cDNA using SuperScript™ III Reverse Transcriptase (Thermo Scientific, 18080093) and Oligo(dT)18 Primers (Thermo Scientific, SO131). Quantitative PCR was performed using a TaqMan™ Gene Expression Master Mix (4369016, Applied Biosystems) *via* StepOne Real-Time PCR Systems (Applied Biosystems). The TaqMan Gene Expression assays used were Hs01005622_m1 (fatty acid synthase, FASN), Hs00168352_m1 (3-hydroxy-3-methylglutaryl-CoA reductase, HMGCR), Hs00996004_m1 (monoglyceride lipase, MGLL), Hs00173425_m1 (lipoprotein lipase, LPL) and Hs00354519_m1 (CD36). The expression of each gene was normalized to the expression of GADPH (Hs02786624_g1). Microarray data were obtained from publicly accessible gene expression profile database GEO (http://www.ncbi.nlm.nih.gov/geo/). Provenzani *et al* [7] performed a microarray study, analyzing differential gene expression in SW480 and SW620 cells (data accessible at NCBI GEO database, accession GSE2509)

### Lipid Extractions

First, the cell pellets were washed with 0.5 mL 0.9% NaCl. For extraction of lipids the pellets were resuspended in 1 ml ice-cold MMC (1:1:1 v/v/v methanol/MTBE/chloroform). Samples were incubated on an ultrasonic bath for two minutes. Phase separation was induced by adding 300 μL MS-grade water. After 10 min incubation, the samples were centrifuged for 10 min at 1000 rpm and the upper (organic) phase was collected. Then 200 μL of collected organic phase were dried in a vacuum rotator and stored at –20 °C until analysis.

### Determination of Total Triglyceride and Cholesterol Ester Contents

Total triglyceride content in the lipid extracts was spectrophotometrically determined using commercially available kit (Analyticon Biotechnologies AG, Catalogue # 5052) against a calibration-curve generated using known concentrations of triglyceride standard (SUPELCO, 17811-1AMP). Cholesteryl esters were quantified using Cholesterol/ Cholesteryl Ester Quantitation Kit (Abcam, ab65359) was used according to manufacturer’s guidelines.

### Lipidomic Profiling and Data Analysis

Dried sample extracts were reconstituted in 100 µL 2:1:1 v/v/v isopropanol/acetonitrile/water. 5 µL aliquots were injected into an ACQUITY I-class ultra-performance liquid chromatography (UPLC) system (Waters, Germany) coupled to an Impact II high-resolution quadrupole time-of-flight mass spectrometer (Bruker Daltonik GmbH, Germany). Chromatographic separation was achieved by gradient elution (%A: 0 min, 60; 1.2 min, 57; 1.26 min, 50; 7.2 min, 46; 7.26 min, 30; 10.8 min, 0; 12.96 min, 0; 13.02 min, 60; 14.4 min, 60) using a buffered solvent system (A: 60:40 v/v acetonitrile/water, B: 90:10 v/v isopropanol:water, both with 10 mM ammonium formate and 0.1% formic acid) and a 2.1 mm × 75 mm × 1.7 µm CSH-C18 column (Waters, Germany) equipped with an 0.2 µm pre-column in-line filter. The flow rate was 0.5 mL min-1 and column temperature was 55°C. Electrospray ionization (ESI) conditions were as follows: polarity (+), capillary voltage, 4500 V, end plate offset, 500 V, nebulizer pressure, 2.5 bar, dry gas (N2) flow, 8 L/min. Ion transfer parameters were set to: Funnel 1 RF, 200 Vpp, Funnel 2 RF, 200 Vpp, Hexapole RF, 50 Vpp, Quadrupole Ion Energy, 5 eV, Low Mass, 100 m/z, Collision Energy, 8.0 eV, Pre Pulse Storage, 6.0 µs, Stepping Mode, Basic, Collision RF, 500-1000 Vpp, Transfer Time, 60-100 µs, Timing, 50/50, Collision Energy, 100-250%. Alternating MS and MS/MS scans were acquired using a Sequential Windowed Acquisition of All Theoretical Fragment Ion Mass Spectra (SWATH) scheme (m/z 350-975, width 25 Da). For internal calibration, Na formate clusters were spiked into the LC effluent at the end of each run. After data acquisition, files were converted to Analysis Base File (ABF) format using a publicly available converter (Reifycs, Japan) and imported into MS-DIAL (Tsugawa et al. 2015). MS-DIAL parameter settings were as follows: Soft Ionization, Data independent MS/MS, Centroid data, Positive ion mode, Lipidomics. Detailed analysis settings were left at default, except for Retention time end (10 min), Alignment Retention time tolerance (0.2 min), Identification Retention time tolerance (3 min) and Identification score cut off (60%).

For each of 6 replicate samples per cell line we calculated within each major lipid class the relative distribution of lipids containing none (SFA0), exactly one (SFA1) and two or more saturated fatty acids (SFA2). To this end, the measured ion intensities for the groups (SFA0, 1 and 2) were summed up and normalized to the total sum of this lipid class. Mean values and standard deviations for the SFA distribution were calculated over the 6 replicates per cell line. Saturation indices were calculated as a ratio of total saturated fatty acid content to total unsaturated fatty acid content [8] within each major lipid class. Total content of saturated fatty acids was calculated by summing up the intensities of all lipids multiplied by the number of saturated fatty acids present within each lipid. Total content of un-saturated fatty acids in each individual lipid class was calculated by summing up the intensities of all lipids multiplied by the number of un-saturated fatty acids present within each lipid. To compare SI over lipid classes we normalized with respect to the SI value obtained for SW480.

### Statistical analysis

The differences between groups were analyzed by ANOVA or t-test (paired or unpaired), where applicable. Statistical analyses and graphical representations for lipidomic data and quantitative RT-PCR data were performed using the R software environment 3.4.2 (http://cran.r-project.org/) or MetaboAnalyst 3.5 (http://www.metaboanalyst.ca/faces/home.xhtml). P-values <0.05 were considered statistically significant and indicated when different.

## Results and Discussion

A previous study has shown a dramatic increase in abundance of total cholesterol esters (CEs) and triglycerides (TGs) in SW620 cells in comparison to SW480 cells [2]. We also assessed the total TG and CE content and observed that SW620 cells display 2.5-3 fold increase in the levels these storage lipids (**supplementary figure 1**). Next, the fatty acid (FA) composition of TGs and CEs was compared. We observed that in SW480 cells the triglycerides (TGs) with ≥2 saturated fatty acids (SFA) were in the highest proportion (i.e. 61%). The proportion of TGs containing 1 or 0 SFA was 35% and 4%, respectively. These proportions were significantly altered in SW620 cells in which the proportion of TGs with ≥2 saturated fatty acids was decreased to 40%, while the proportions of TGs containing 1 or 0 SFA were respectively increased to 53% and 7%. In agreement to these results, the TG saturation index (SI) was markedly higher in SW480 cells than in SW620 cells. On the other hand, FA-composition and SI of diacylglycerols (DGs), the direct precursors to TGs, was only slightly altered. The FA-composition of CEs also showed minor changes, with slight shift in the proportions of CEs containing 0 or 1 SFA. However, the SI of CEs was also significantly decreased in SW620 cells. We also compared the FA composition of the most abundant membrane lipids, phosphatidylcholines (PCs) and phosphatidylethanolamine (PEs), in SW480 and SW620 cells. No substantial differences were observed in fatty acid composition and SI of PCs and PEs (**Figure 1**). Hence, TGs constitute the major lipid class that displays changes in FA composition in SW620 cells when compared to SW480 cells.

**Figure 1:**
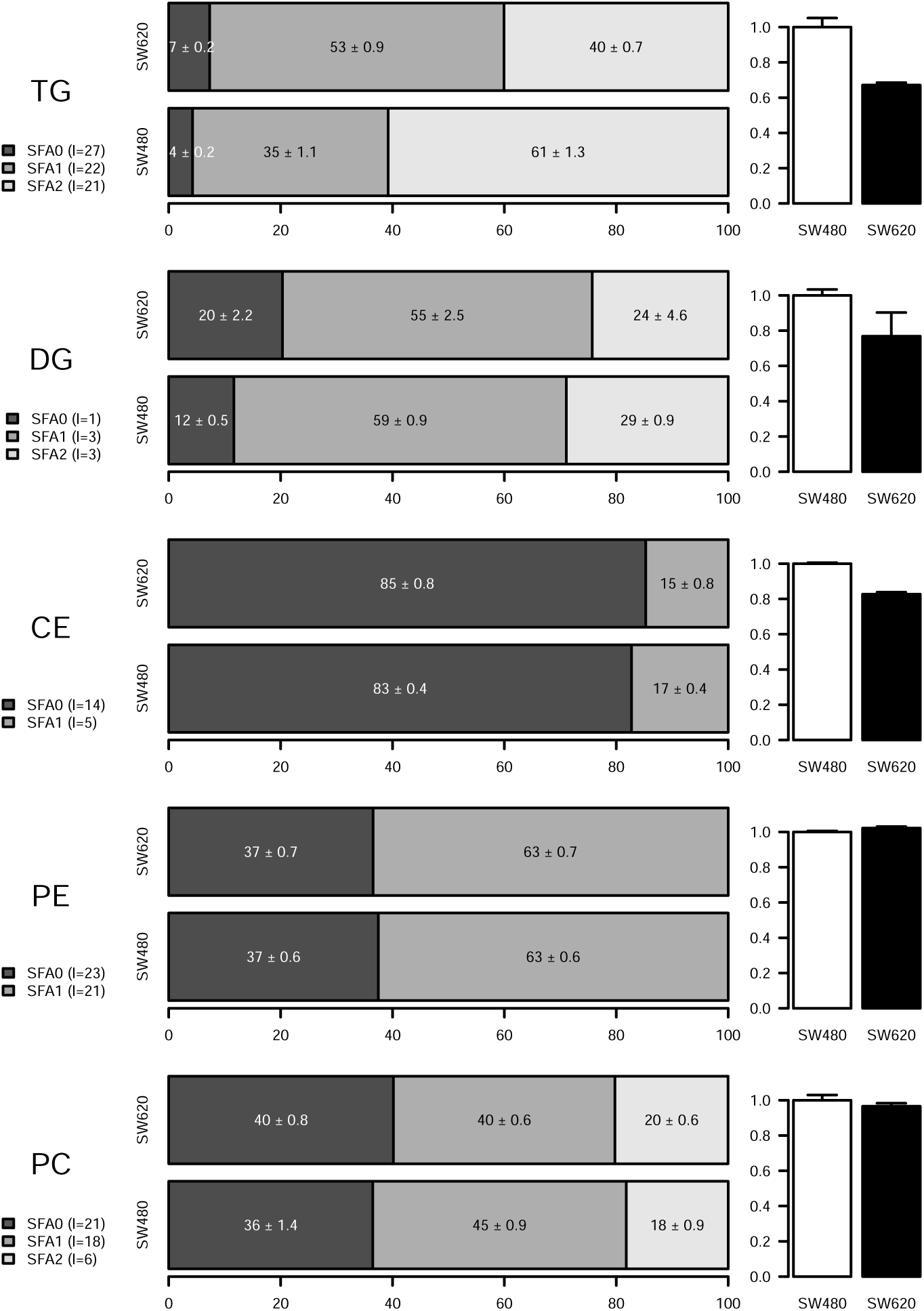
Comparison of fatty acid composition and saturation indices in major lipid classes in SW480 and SW620 cells. The figure is structured in five rows and three columns. Rows show data for the major lipid classes Triglycerides (TG), Diglycerides (DG), Cholesterol Esters (CE), Phosphatidylethanolamines (PE) and Phosphatidylcholines (PC), Left column displays lipid class ID and a legend indicating the number of lipid-subspecies within each lipid class containing different numbers of saturated fatty acids (0, 1 or ≥2SFAs). Center column shows the relative amount (in per cent of summed ion intensity) of lipid subspecies within both SW cell lines. Numeric values for mean and standard deviation calculated over 6 replicate samples per cell line are given within the bars. Right column displays the saturation indices (cf. Method section) of the respective lipid class normalized to SW480.

To understand the molecular mechanisms regulating these changes in the lipidomic profiles, we performed expression analyses and also used publicly accessible gene expression profile database GEO (http://www.ncbi.nlm.nih.gov/geo/). **Figure 2** displays overview of the major cancer-associated lipid metabolism pathways. The fold-changes in the expression levels of various genes/enzymes are indicated with a colored-box on *left-side* of the respective gene symbol. For color-codes see color-key on *top-left* corner of **Figure 2**. As discussed above, the differences in the FA composition of TGs was the most significant difference between lipidomic profiles of SW480 and SW620 cells, with SW620 cells displaying significantly lower TG saturation-index. A recent study has shown that gene silencing of diglyceride acyltransferases (DGATs) –enzymes that catalyze formation of TGs from DGs and Acyl CoA– induces increase in TG saturation-index [9]. Hence, increased expression of DGAT might induce decreased SI in TGs. Our analyses revealed that expression of DGAT was slightly decreased (1-1.5 Fold) in SW620 cells (**Figure 2, TG Synthesis**). Hence, DGAT expression might not be responsible for the changes in TG saturation-index of SW620 cells.

**Figure 2:**
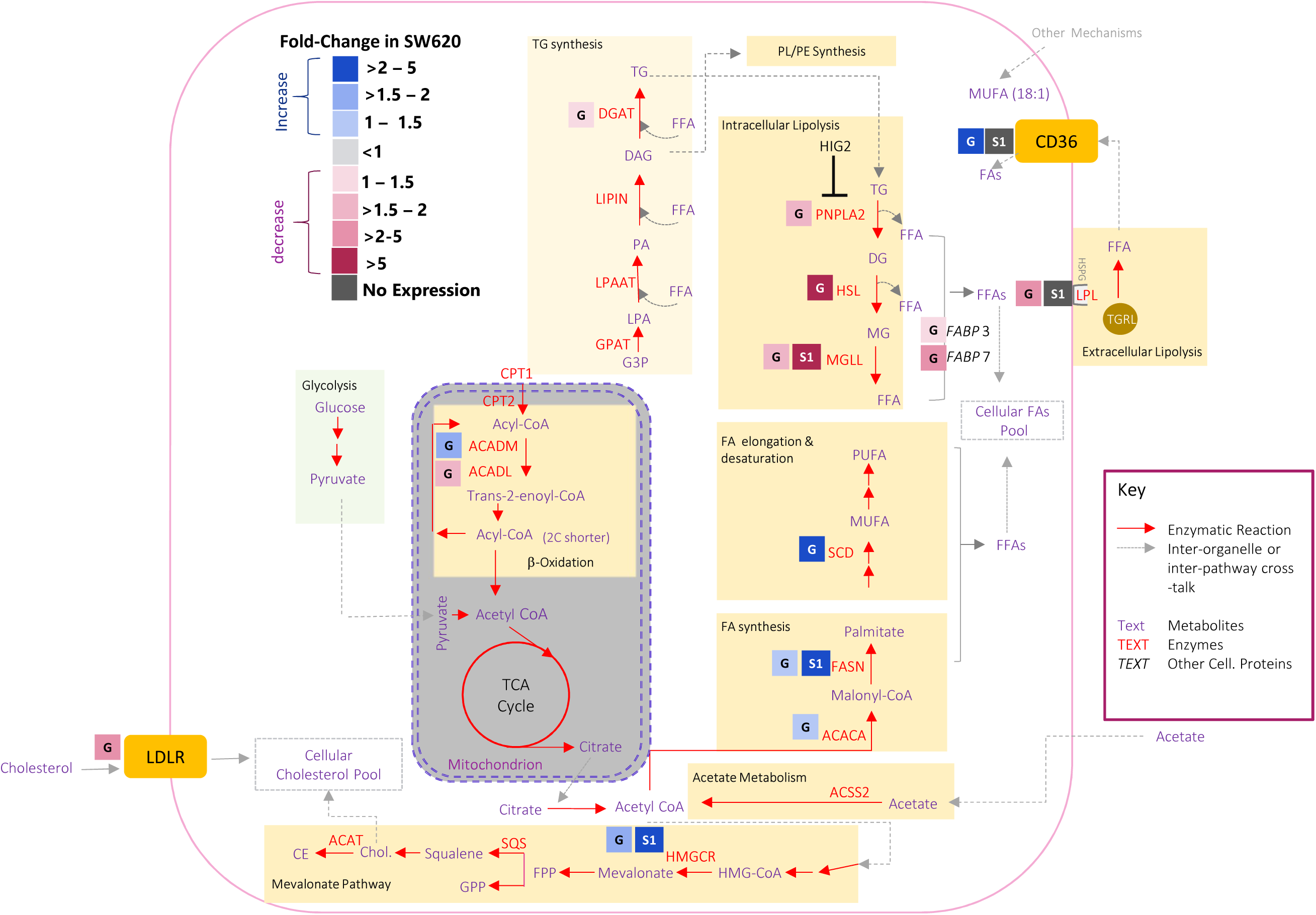
Fold-changes in expression levels of various genes in major lipid metabolism pathways in SW620 *vs.* SW480 cells. The figure highlights all the key lipid metabolism pathways activated in cancer cells. Major pathways are shown as boxes without outlines. The systematic names of these pathways are given in each box. The fold-changes in the expression levels of various genes/enzymes in SW620 cells compared with SW480 cells are indicated with a colored-box on *left-side* of the respective gene symbol. For color-codes see color-key on *top-left* corner. The expression data for these analyses were extracted from at NCBI GEO database, accession GSE2509. For some markers gene expression analysis was performed (**Supplementary Figure S2**). The symbols within fold-change boxes represent the data source (G; GEO database, S, Supplementary Figure S2). **Abbreviations:** ACACA, acetyl-CoA carboxylase 1; ACACB, acetyl-CoA carboxylase 2; ACSS2, acyl Co-A synthetase-2; ADFP, adipose differentiation protein; ATGL, adipose triglyceride lipase; FA, fatty acids; FABP3, fatty acid binding protein 3; FABP7, fatty acid binding protein 7; FFA, free fatty acids; FASN, fatty acid synthase; H, hypoxia; HIF-1α, hypoxia-inducible factor 1-alpha; HIF-2α, hypoxia-inducible factor 2α; HIG2, hypoxia-inducible gene 2 protein; HMGCR, 3-hydroxy-3-methylglutaryl-CoA reductase; LD, lipid droplet; LCAD, long-Chain Specific acyl-CoA dehydrogenase; MAG, monoacylglycerol; MCAD, Medium-chain acyl-CoA dehydrogenase; MUFA, monounsaturated fatty acids; PBMCs, peripheral blood mononuclear cells; PC, Phosphatidylcholines; PE, phosphatidylethanolamines; Pcho, propargyl-choline; PI, phosphatidylinositol; PS, phosphatidylserine; PUFA, polyunsaturated fatty acids; SCD, stearoyl-CoA Desaturase; SFA, saturated fatty acids; SREBP, sterol regulatory element-binding proteins; TG, triglycerides.

Next, we looked at other lipid metabolism pathways that have been implicated in regulation of lipid saturation-indices. Highly active *de novo* FA synthesis induces increased SFAs or MUFAs levels in cancer cell cells [10]. The expression of FA synthesis genes is up-regulated in SW620 cells, hence this pathway is also not contributing in decreased SI in these cells (**Figure 2, FA Synthesis**).

Major genes of intracellular lipolytic pathway –including PNPLA2, HSL and MGLL– are all down regulated in SW620 cells (**Figure 2, Intracellular Lipolysis**). This may explain the dramatic increase in total triglyceride (TG) levels in SW620 cells. However, specific decrease in ≥2 SFA containing TGs could not be explained by this mechanism.

The expression of ACADM and ACADL –the enzymes that catalyze the first step of FA oxidation in mitochondria– is differentially regulated in SW620 cells. The expression of ACADM was up regulated, while the expression of ACADL was down-regulated in SW620 in comparison to SW480 cells (**Figure 2, FA Oxidation**). This observation is of special interest in this context, because ACADL is known to mediate unsaturated FA oxidation, whereas ACDM prefers saturated FAs as substrates. Hence, it is possible that in SW620 cells there is an increased oxidation of SFAs, while decreased oxidation of unsaturated FAs. This may cause selective depletion of SFAs in the stored lipids and induced decreased SI of TGs and CEs. Recent studies have shown that disruptions in lipid metabolism or nutrient/oxygen supply also drastically affects FA-composition of TGs in cancer cells, while the FA-composition of other lipid classes is not significantly affected [9,11]. Hence, FA-composition of TGs is most markedly affected by the changes in cancer stage, tumor microenvironment and de novo FA synthesis. Further studies are required to understand the significance of fatty acid composition of TGs in cancer progression and survival.

## Abbreviations

FASN: fatty acid synthase
HMGCR: 3-hydroxy-3-methylglutaryl-CoA reductase
MGLL: monoacylglycerol lipase
LPL: lipoprotein lipase
PC: Phosphatidylcholines
PE: Phosphatidylethanolamines
PPE: Plasmenylphosphatidylethanolamines
PPC: Plasmenylphosphatidylcholines
TG: Triacylglycerols
CE: Cholesterol esters and DG Diacylglycerols.

## Declarations

### Ethics approval and consent to participate

Not applicable.

### Consent for publication

Not applicable.

### Availability of Data and Material

The datasets supporting the conclusions of this article are included within the article and supplementary files.

## Acknowledgments

We are grateful to Mr. Banaras Masih for his technical assistance for this project.

## Authors’ Contributions

This study was designed by NZ. Experimental work was done by NZ, CJ and RM; and data was analyzed and interpreted by JL, CJ, NZ and RM. The manuscript was written by RM and NZ; and reviewed and approved by all authors.

---

**Supplementary Figure S1:**
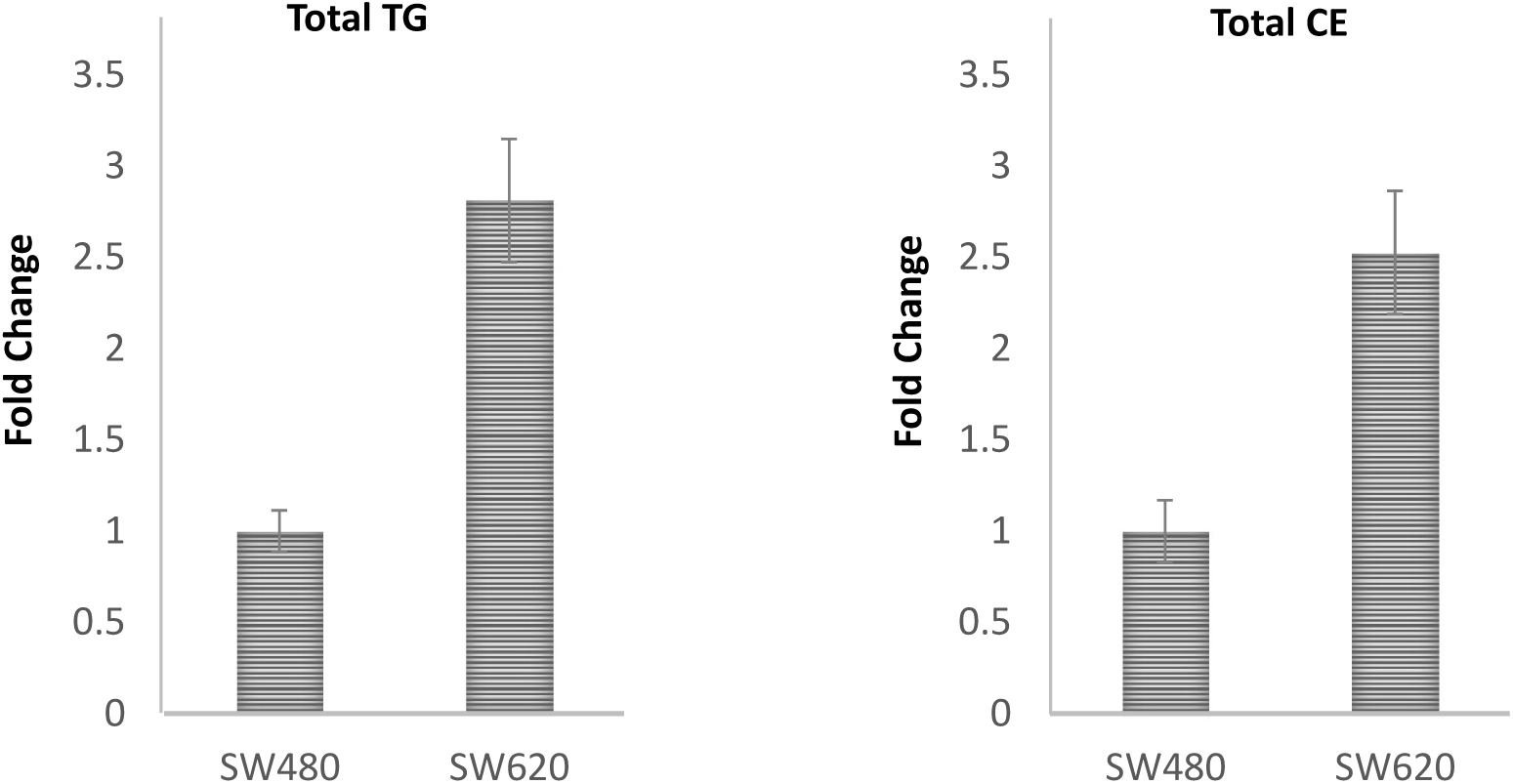
Fold-changes in total triglyceride(TG) and cholesterol ester (CE) content in SW480 vs. SW620 cells.

**Supplementary Figure S2:**
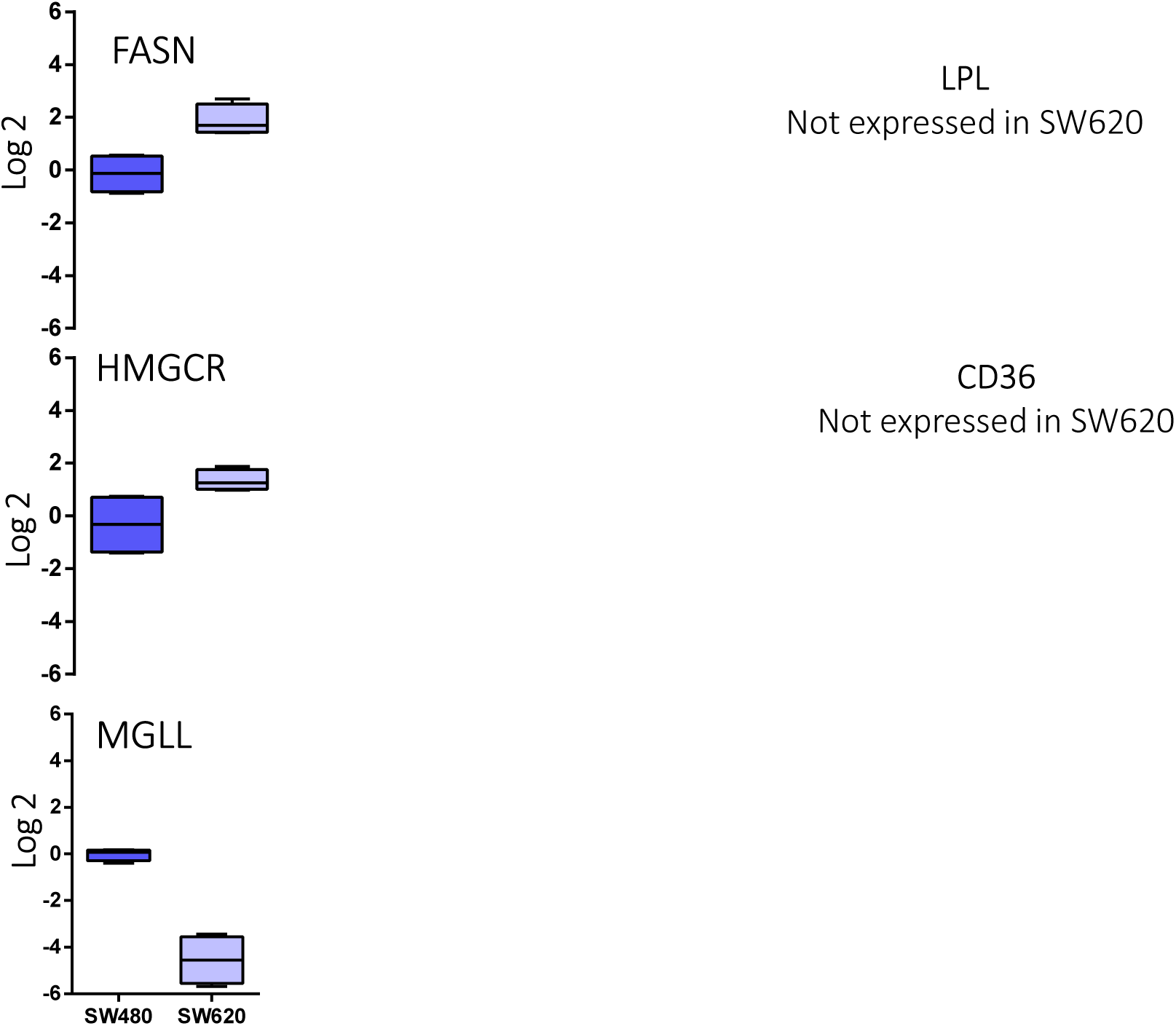
Expression of selected genes from de novo lipid synthesis or lipid uptake/degradation pathways in SW480 and SW620 cell lines. Box plots showing log2 transformed and median normalized values of FASN, HMGCR and MGLL. LPL and CD36 expressions were not detected. The levels of the different transcripts were measured in 3 to 6 samples by qPCR. The results show the distribution of corresponding transcripts relative to GAPDH, with the box indicating the 25th–75th percentiles, with the median indicated. line. The whiskers show the range.

